# Fins as a reliable surrogate tissue for age-related changes of telomeres and DNA methylation in gonads of a short-lived fish

**DOI:** 10.1101/2025.08.29.672996

**Authors:** Milan Vrtílek, Anna Kromerová, Malahat Dianat, Miloslava Fojtová, Dagmar Čížková, Jiří Fajkus

## Abstract

Senescence is a multifactorial and individualised process of age-related physiological decline. Cellular markers, such as telomere length and DNA methylation, can reveal subtle changes associated with chronological age or expected lifespan. In this study, we evaluated the utility of fin tissue as a surrogate for assessing telomere length and proportion of DNA methylation in the gonads of a small, short-lived laboratory fish, the turquoise killifish (*Nothobranchius furzeri*). We collected fin and gonadal tissues from both females and males at three different ages, and extracted DNA to measure telomere length via terminal restriction fragment (TRF) analysis and global DNA methylation levels using double-digest restriction-associated DNA sequencing (ddRADseq). Our results show a notable correspondence between telomere length and DNA methylation patterns in fin and gonadal tissues. These findings support the use of fin biopsies as a non-lethal method for assessing ageing biomarkers in the gonads of small freshwater fish.

## INTRODUCTION

Senescence is an age-related process of bodily decline expressed as dysbalanced physiology, reduced reproduction and lower survival (Jones et al., 2014). The onset and rate of senescence typically vary between individuals within the same population (Bryant and Reznick, 2004; Patrick et al., 2022). Focusing on cellular markers of senescence obtained through non-lethal sampling of an individual may provide important information on age-related decline across different tissues and before more visible changes are evident (Horvath and Raj, 2018; López-Otín et al., 2013).

Telomeres are nucleoprotein complexes at the ends of eukaryotic chromosomes, ensuring their integrity (Blackburn et al., 2015). Telomeres inevitably become shorter with each cell division and are additionally damaged by various factors such as environmental pollution or stress, including reactive oxygen species (Armstrong and Boonekamp, 2023; Monaghan, 2014). Consequently, shorter telomeres are typically found in older individuals across most vertebrates (Remot et al., 2022). Telomere attrition is therefore considered one of the canonical markers of cellular senescence (López-Otín et al., 2013). While distinct telomere shortening may trigger cell apoptosis, it is partially counteracted by an enzyme that extends telomeres - telomerase. Telomerase is particularly active in stem cells and highly proliferative tissue such as germline (Blackburn et al., 2015; Harel et al., 2015; Pech et al., 2015). In contrast, telomerase activity is suppressed across somatic tissues in birds and mammals (Gomes et al., 2010). This may lead to uncoupling of the age-related dynamics between somatic and germline telomeres (Ramos-Ibeas et al., 2019). In ectotherms, however, high telomerase activity is retained in somatic tissues (Anchelin et al., 2011; Gomes et al., 2010). Very few studies, and with contrasting results, on the relationship between somatic and gonad or germline telomere lengths is currently available from ectotherms (Carneiro et al., 2016; Lunghi and Bilandžija, 2024; Morbiato et al., 2023).

DNA methylation, the addition of a methyl group to the cytosine base, has recently been proven as a reliable chronological marker for calibrating species-specific “epigenetic clock” in multiple vertebrates, including fish (Anastasiadi and Piferrer, 2020; Horvath and Raj, 2018; Lu et al., 2023; Mayne et al., 2021). DNA methylation is not a completely neutral marker, however. Methylation of so-called CpG islands (sites with high density of cytosine-guanine doublets) close to gene promoters is one of the main mechanisms regulating gene expression (Keshet et al., 1985). While the overall level of DNA methylation tends to decrease with age, methylation of CpG islands increases (Jones et al., 2015). The wide usage of epigenetic clock underscores that the age-related DNA methylation patterns are often shared across different tissues (Giannuzzi et al., 2024; Horvath and Raj, 2018; Le Clercq et al., 2023). Although DNA methylation allows precise calibration for specific purposes, little is known about the shared patterns between somatic and gonad tissues. Since these tissues can show distinct aging dynamics, such differences might limit versatility of age-related methylation patterns.

Telomere length and DNA methylation represent two largely independent and complementary age-related markers (Sheldon et al., 2022). Telomere length is more associated with what can be called the biological age or individual quality (Angelier et al., 2019), and is a relatively good predictor of the lifespan within a population (Bize et al., 2009; Heidinger et al., 2012). Telomeres are, on the other hand, poor predictors of individual chronological age, especially when compared to the emergent properties of DNA methylation (Le Clercq et al., 2023). The multivariate character of DNA methylation data allows calibration to estimate either real (chronological) age or biological age (i.e., expected lifespan) (Giannuzzi et al., 2024).

Here we compare age-related patterns in telomere length and DNA methylation between fins and gonads in a small freshwater fish, turquoise killifish (*Nothobranchius furzeri* Jubb 1971). Turquoise killifish is a short-lived fish adapted to seasonal ponds in the African savannah with captive lifespan typically shorter than one year (Cellerino et al., 2016). The short lifespan makes the turquoise killifish an attractive vertebrate model species for studying senescence (Brunet, 2020; Poeschla and Valenzano, 2020; Reichard and Polačik, 2019). Telomeres shorten with age in somatic tissues of both females and males (Hartmann et al., 2009), although females exhibit longer telomeres in their fins at a given age compared to males (Reichard et al., 2022). Telomerase expression is particularly high in the gonads of turquoise killifish males (Harel et al., 2015). Recently, age-related DNA methylation in fins and brain of the turquoise killifish was thoroughly studied to develop an epigenetic clock (Giannuzzi et al., 2024). It seems that the ageing dynamics in DNA methylation patterns of these two tissues are relatively well aligned, allowing for identification of shared batteries of methylation marks (Giannuzzi et al., 2024).

In the present study, we tested the potential utility of fin tissue as a surrogate for age-related telomere attrition and DNA methylation alterations in the gonads of female and male turquoise killifish. We chose gonads due to their relatively high proliferation rate and their role in potential transmission of age-related patterns to the next generation. Our study is inevitably cross-sectional, with the future objective to sample only fins without the need to sacrifice the studied animal to obtain gonads. This is especially topical for small laboratory fish, where fin biopsies allow for repeated collection of tissue material (unlike blood, for example). Our main aim was to test the correlation between fin and gonads, both for telomere length and proportion of DNA methylation. Based on previously reported data, we predicted shorter telomeres across tissues in older individuals (Hartmann et al., 2009), and shorter telomeres in males compared to females at a specific age (Reichard et al., 2022). Finally, we expected that methylation of CpG islands will increase with age.

## METHODS

### Overview

We designed a common garden cross-sectional study to collect samples of fin and gonad from individually-housed females and males of the turquoise killifish at three prespecified time-points covering most of the population’s expected lifespan. The DNA isolated from these samples was used to measure the telomere length and to generate libraries for the analysis of DNA methylation.

### Fish husbandry

We worked with a wild-derived strain of the turquoise killifish (MZCS 222 *Nothobranchius furzeri* Jubb 1971). This strain was maintained at the fish breeding facility of the Institute of Vertebrate Biology, CAS in Brno, Czech Republic, since its import in 2011 (originating from 20 males and 40 females) (Cellerino et al., 2016). Fish for this study came from clutches collected from group spawning of 5 stock tanks with a total of 75 fish. The clutches were incubated on peat in a zip-lock bag at 18.5 °C for 45 months. Before hatching, we collected and sterilized the eggs with the solution of peracetic acid (0.5 mL/L of the Persteril 5, FICHEMA) in 3 cycles (5 min in the sterilization solution and 5 min in reverse osmosis - RO - water). We stored them on moist filtration paper in a sealed Petri dish at 27 °C for three weeks to stimulate completion of embryonic development.

The experimental fish were hatched by watering the eggs with aged tap water (22 °C, 0.450 mS/cm) in 2-L containers. Early on, fish were fed with *Artemia sp. nauplii* for 8 days and then weaned on bloodworms (adult diet). Husbandry followed the published protocol (Polačik et al., 2016). Eight days after hatching, the fish were distributed into individual 3.5-L tanks in common-garden conditions of a recirculation system with 27 °C, 1 mS (mixed from RO water and NaCl), and 14:10 light:dark regime. All fish were sexually dimorphic by day 23 post-hatching, and we began with regular morning spawning at 35 days (5 weeks). We spawned the fish twice a week in system water in 2-L container on top of a black 5-mm mesh for 1.5 h. We thus kept each individual in standardized isolated conditions while allowing for regular reproduction. The experiment was finished at the last sampling point, at the age of 30 weeks (8.5 months) when last surviving fish were sacrificed.

### Tissue sampling and DNA isolation

We collected fin and gonad tissues from the experimental fish at three pre-specified time points (referred to as AGE1, AGE2, and AGE3; Fig. 1). The timing of the sampling was based on the female survival curve in this strain from a previous study (Žák and Reichard, 2021), individual males live longer. The three points correspond with full female maturity (early adulthood, all females have ovulated eggs, 8 weeks, AGE1), 50% (prime age, peak of egg production, 22 weeks, AGE2) and 25% (senescence, reproductive decline, 30 weeks, AGE3) survival of females in the population, respectively. Our aim was to capture age-related variation in the cellular markers between the two selected tissues – fin and gonad. Longitudinal studies are pivotal in senescence research, and we would thus like to provide a platform for future experiments with turquoise killifish or similar small fish. We simulated repeated sampling of the same individual while avoiding regenerated tissue by collecting different parts of the fins (similar to (Baumgart et al., 2016) at AGE1 (upper half of caudal fin), AGE2 (lower part of caudal fin) and AGE3 (dorsal fin) (Fig. 1).

**Figure 1.**
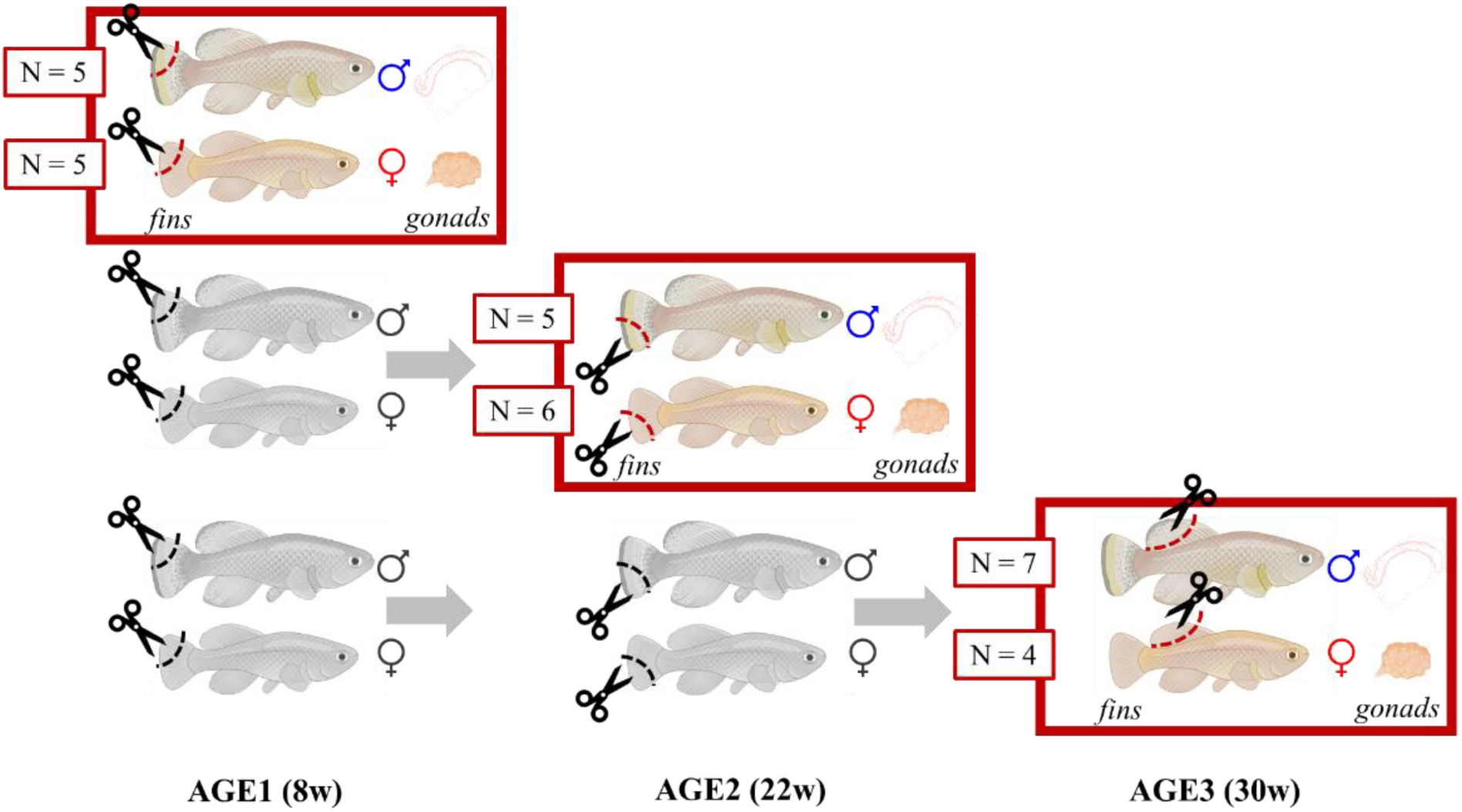
The cross-sectional design of sampling fins and gonads from turquoise killifish. We sacrificed both females and males at three time-points (red squares). We collected upper half of caudal fin at AGE1 (8 weeks), lower part of caudal fin at AGE2 (22 weeks) and dorsal fin at the final AGE3 (30 weeks), always together with the gonads (ovaries in females or testes in males). In AGE1 and AGE2, we also cut fins of the remaining live fish (in gray). Designed using BioRender.

We randomly selected females and males for sacrificing at AGE1 (5+5), at AGE2 (6+5), and then took the survivors (4+7) at the last AGE3 sampling. The random selection was performed using *sample* function in R software (ver. 4.4.1; R Core Team, 2024). Sampling was always carried out between 8 and 11 AM, so that the fish had been fasted for at least 16 hours. Euthanasia was performed using a clove oil overdose. We measured body length and weight, dissected body weight and gonad weight (Table S1). Fin tissue was trimmed and whole gonads were also collected and these samples were stored at -80 °C for DNA isolation. To maximise DNA yield, all ovulated and maturing oocytes were removed from the ovaries prior to processing.

DNA was extracted from the turquoise killifish fin and gonad using the cetyltrimethylammonium bromide method (Kovařík et al., 2000). The DNA integrity was verified by standard horizontal agarose electrophoresis, and DNA concentration was measured by the NanoDrop spectrophotometer (Thermo Fisher Scientific) and Qubit (Thermo Fisher Scientific). The extracted DNA was aliquoted and used for both telomere and methylation analyses.

### Telomere length analysis

Telomere length was assessed by the terminal restriction fragment (TRF) method as described in (Fojtová et al., 2015). Five μg of genomic DNA was digested by the HinfI and RsaI restriction endonucleases (NEB) and separated with standard agarose gel electrophoresis. Samples were transferred by Southern blot to the positively charged nylon membrane (ROTI®Nylon plus), and hybridized with a radioactively labelled telomere probe, which was synthesized by non-template PCR (Adamusová et al., 2020; IJdo et al., 1991). Telomere-specific hybridisation signals were visualised by phosphoimager FLA 7000 (FujiFilm) and evaluated by the WALTER online toolset (Lyčka et al., 2021).

### ddRAD sequencing

For preparation of the double-digest restriction site-associated DNA (ddRAD) libraries from the 64 samples, we combined the traditional approach of Peterson et al. (2012) and a 2RAD approach by Bayona-Vásquez et al. (2019) as described by Komarova et al. (2021) (for details see Appendix 1). Briefly, each DNA sample was digested with two enzyme pairs: a common cutter that is insensitive to cytosine methylation (MspI) and a rare cutter (EcoRI), and alternatively with the methylation-sensitive common cutter (HpaII) and EcoRI. Thus, two libraries per sample were generated. One library with all CCGG restriction sites cut (MspI common cutter used; “RAD library”), and the other where methylated sites remain uncut (HpaII common cutter used; “EpiRAD library”). After digestion, adapters were ligated and DNA fragments were purified using SpriSelect beads (Beckman Coulter Life Sciences) (1× volume ratio). The fragments were amplified using dual-indexed primers, and libraries were checked via electrophoresis. All libraries were pooled in equimolar amounts and concentrated with SpriSelect beads (1.2×). The pooled library was size-selected in tight mode, with average fragment size of 340 base pairs (bp) using Pippin Prep (Sage Science). The final library was sequenced on the Illumina NovaSeq X Plus PE150 platform (Novogene Co. Ltd.), generating 150 bp paired-end reads.

We sequenced 128 libraries in total. The quality of the raw sequencing reads was assessed using FastQC (ver. 0.11.9; Andrews, 2010) and summarized with MultiQC (ver. 1.8; Ewels et al., 2016). Inline barcodes, residual adapters, restriction enzyme recognition sites, and low-quality bases were trimmed using Skewer (ver. 0.2.2; Jiang et al., 2014). After trimming, the average number of reads per sample was 3.76 million for RAD and 4.45 million for EpiRAD libraries. The assembly was performed using IPYRAD (ver. 0.9.92; Eaton and Overcast, 2020) with the *N. furzeri* MZM-0403 reference genome (GCF_027789165.1). Refer to the Appendix 2 for the parameters’ setting used in the assembly.

First, we generated a comprehensive set of all reference ddRAD loci. The dataset was assembled by concatenating RAD and EpiRAD libraries for each sample, to minimize the potential loss of loci with low depth in individual libraries. The final number of assembled loci across concatenated libraries was 37 902 (19 249 to 26 118 loci per library, with a mean of 23 345 loci). Read counts for the reference loci in each RAD and EpiRAD library were then extracted. We used SAMtools (ver. 1.14; Danecek et al., 2021) with the *coverage* command to calculate per-locus coverage statistics for each individual.

### Identification of methylated loci

Before the analysis of read counts and loci filtering, libraries were standardized to a common size. Counts per million (CPM) standardization was applied to obtain relative read representation of each locus in a library, where the number of reads per locus was divided by the total number of reads mapped to the reference loci (library size) and then multiplied by one million. Loci insufficiently represented across the RAD (methylation non-sensitive) libraries were then filtered out. After the CPM standardisation, loci with at least 15 reads in the RAD libraries across all 64 samples were retained for analysis (Figure S1, Table S2). This threshold removed non-conclusive loci for assessment of methylation status while keeping a sufficient number of loci for subsequent differential methylation analysis. The resulting reduced dataset contained 7 841 loci which were used to compare the proportion of methylated loci across samples. Almost identical results were obtained when alternative thresholds of 7 or 21 were applied (Table S2).

A locus was defined as methylated when no reads were detected in the EpiRAD (methylation sensitive) library. This approach may lead to an underestimation of overall DNA methylation within the samples, as loci that are partially methylated (e,g., hemi-methylated or non-uniformly methylated across sampled cells) were classified as non-methylated. At the tissue level, DNA methylation of a locus represents a continuous variable; however, accommodating this would require the introduction of a subjective cut-off, thereby increasing analytical ambiguity (Dimond et al., 2017).

### Data analysis

All data analyses were performed using R software (ver. 4.4.1; R Core Team, 2024).

To compare the length of telomeres or proportion of DNA methylation between fins and gonads, we used simple two-tailed t-test. The effects of SEX, AGE and total body size on median telomere length in fins or gonads were analysed using general linear models. An empty (intercept only) model was compared with models including additive effects of the explanatory variables, as well as with the most complex model incorporating the SEX×AGE interaction together with the additive effect of body size. Model selection was based on Akaike Information Criterion for small sample sizes (*AICc*), implemented in the package MuMIn (ver. 1.48.11; Bartoń, 2025) and the model with lowest AIC was chosen for interpretation. Model-estimated means with 95% confidence intervals were used for visualisation in plots. The goodness of fit for correlations in telomere length or proportion of DNA methylation between fin and gonad tissue was quantified using the model coefficient of determination (R^2^; Nakagawa et al., 2017).

Loci uniformly methylated (1) or non-methylated (0) across samples were identified by sub-setting the data (e.g., female fins of AGE1) and calculating row sums. For clustering analysis, loci that were uniformly methylated or unmethylated across all samples were excluded, and the focus was placed on differentially methylated loci. Dissimilarity in DNA methylation states between samples was then assessed using classical multidimensional scaling ordination (function *cmdscale*, package ape ver. 5.8; Paradis and Schliep, 2019).

## RESULTS

Our cross-sectional population sample consisted of fin and gonad tissue collected from 32 individuals (17 males and 15 females, Table S1) across three age groups to analyse telomere length and DNA methylation.

### Effect of age on median telomere length in fins and gonads

Substantial variation in median telomere length was observed among individual fish. The telomere length median ranged between 4.38 and 9.24 kb in fins, and 3.15-9.92 kb in gonads (Figure S2). Telomere length was comparable between fins and gonads (fins: 7.29 ± 0.20 kb, gonads: 6.85 ± 0.29 kb [mean ± s.e.]; t-test: t54.5 = 1.25, P = 0.215).

Telomeres were not shorter in the fish sampled at older age. Interestingly, fish from the middle age sampling (AGE2, 22 weeks) had shorter telomeres in fins than individuals from the younger AGE1 (8 weeks) or older AGE3 (30 weeks) samplings (effect of AGE: F2,28 = 8.50, P = 0.001; Fig. 2A). In all groups tested, males tended to have shorter telomeres than females in both fins (F1,28 = 3.42, P = 0.075, Fig. 2A) and more clearly in gonads (F1,28 = 43.71, P < 0.001, Fig. 2B). The AGE-group differences in telomere lengths were more pronounced in male (in contrast to female) gonads (AGE×SEX interaction: F2,28 = 5.30, P = 0.012; Fig. 2B).

**Figure 2.**
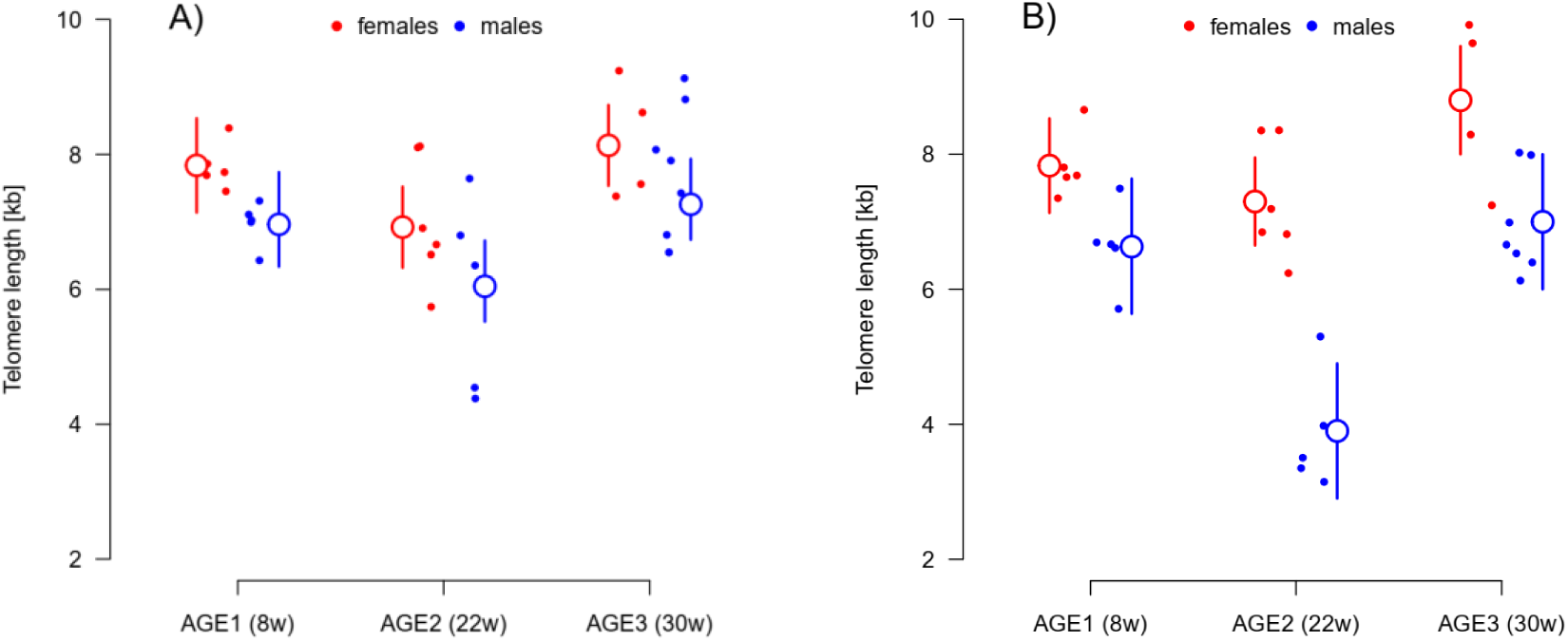
Medians of telomere lengths at the three AGE-samplings in fins (A) and gonads (B). Males (in blue) had generally shorter telomeres than females (in red), although this difference was only significant in gonads (i.e., shorter telomeres were recorded in testes than ovaries). AGE1 group was sampled at 8 weeks post-hatching, AGE2 at 22 weeks, and AGE3 at 30 weeks. The small full points show raw values of telomere length median for each individual. The larger empty points show mean estimate for respective AGE-group and sex separately, with the vertical lines corresponding to the 95% confidence interval (CI). For the distribution of telomere-specific hybridization signals in individual samples see Figure S2.

### Correlation between telomere length in fins and gonads

Telomere length in fins was predictive of telomere length in gonads. The correlation between telomere lengths in fins and gonads in the 32 individuals studied was strong (adjusted R^2^ = 0.733) and positive (slope: 0.97 ± 0.14, t1,29 = 6.805, P < 0.001; Fig. 3). Females and males differed in the intercept of the slope. Females had relatively longer telomeres in gonads for given telomere length in fins compared to males (the additive male effect: -1.36 ± 0.31, t1,29 = -4.35, P < 0.001).

**Figure 3.**
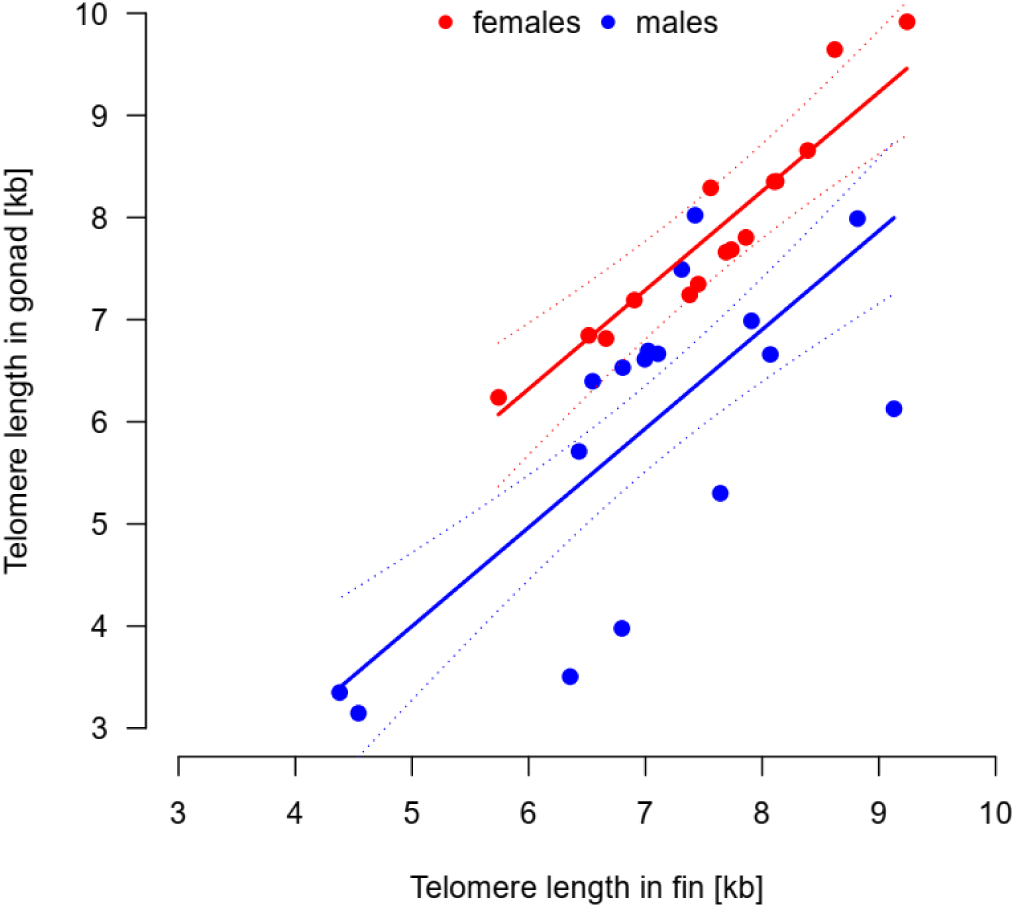
Medians of telomere length in fins and gonads are positively correlated in both females and males. The solid trendlines illustrate the additive effect of sex - females (red line) have relatively longer telomeres in gonads than males (blue line) when controlling for the telomere length in fins. The dotted lines give upper and lower limits for 95% confidence intervals of the trendline estimates. The points show medians of telomere length per individual, red points for females and blue points for males.

### Effect of age on the proportion of methylated DNA loci in fins and gonads

We identified 7 841 well-represented loci in our dataset (i.e., loci with at least 15 reads across all methylation-non-sensitive RAD libraries after the CPM standardization). We then classified DNA methylation in the samples according to the number of reads in a locus in the methylation-sensitive EpiRAD library – loci with zero reads in the EpiRAD library were considered as methylated and non-zero as unmethylated. Based on this approach, the proportion of methylated loci across individual samples varied between 0.47 and 2.17% in fins and 0.38-5.97% in gonads. Fins had on average a lower proportion of methylated loci than gonads (fins: 1.36 ± 0.10%, gonads: 2.15 ± 0.24% [mean±s.e.]; t-test: t40.7 = 1.01, P = 0.004).

The proportion of methylated loci increased with age. In fins of young adults (AGE1), a lower proportion of methylated loci was detected compared to old fish (AGE3), irrespective of their sex (effect of AGE: F2,28 = 4.57, P = 0.019; effect of SEX: F1,28 = 0.21, P = 0.644; Fig. 4A). In gonads, the differences between AGE-groups were not significant (effect of AGE: F2,28 = 2.88, P = 0.073), male gonads showed a higher proportion of DNA methylation than females across age groups (F1,28 = 8.18, P = 0.008; Fig. 4B).

**Figure 4.**
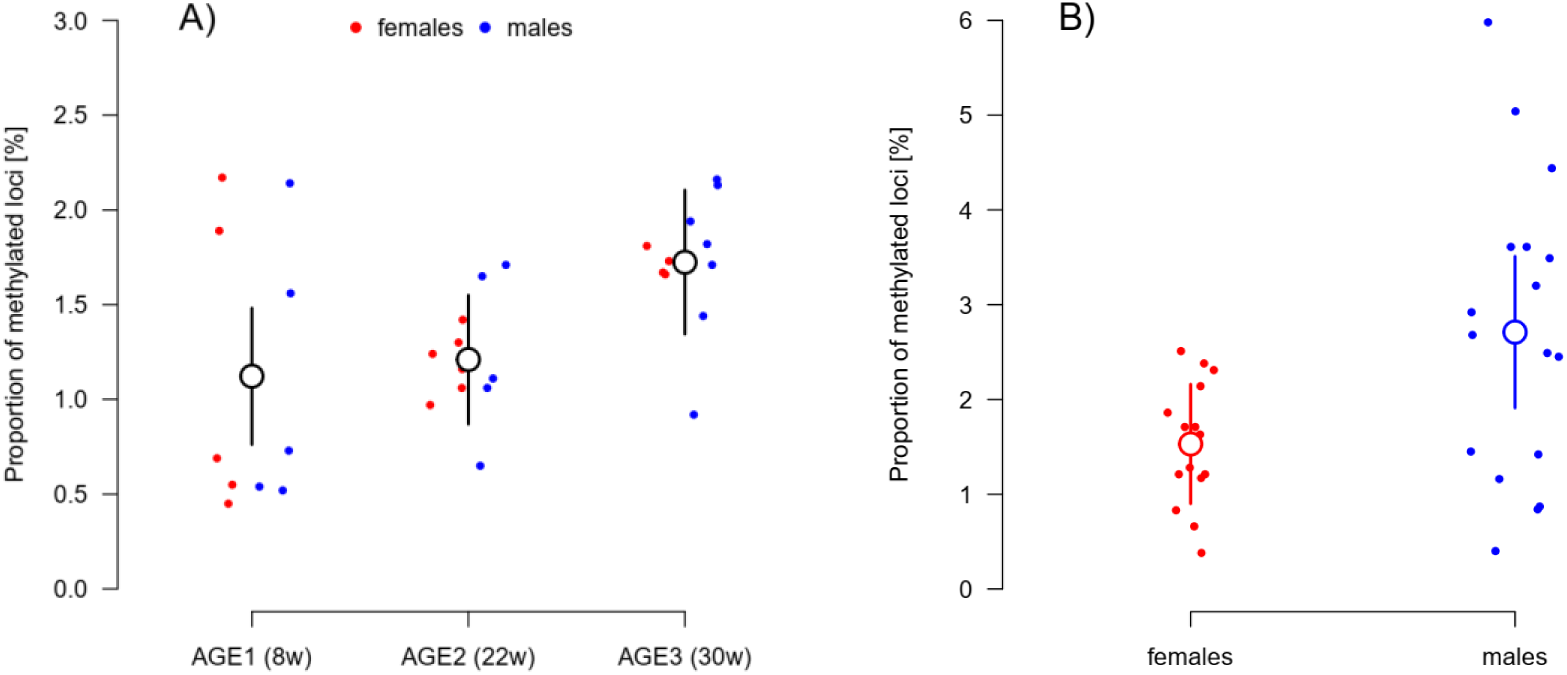
The proportion of methylated loci in fins (A) and gonads (B). In fins, there was an apparent difference between the AGE-groups (larger empty black points) but not between females (red points) and males (blue). In the gonads, only the sex difference was pronounced (testes in blue vs. ovaries in red). The larger empty points show mean estimates for the corresponding AGE-group (A, fins) and SEX-group (B, gonads), respectively. The vertical lines show the 95% CI of these estimates.

### Relationship between proportion of methylated loci in fins and gonads

The proportion of DNA methylation in fin and gonad sampled from the same individual corresponded well (adjusted R^2^ = 0.689). The overall relationship was positive, but the slope was sex-specific (DNAme.fin×SEX interaction: F1,28 = 5.67, P = 0.024). The fin-gonad slope of the proportion of methylated loci was steeper in males compared to females (male-specific slope: 1.21 ± 0.51; Fig. 5).

**Figure 5.**
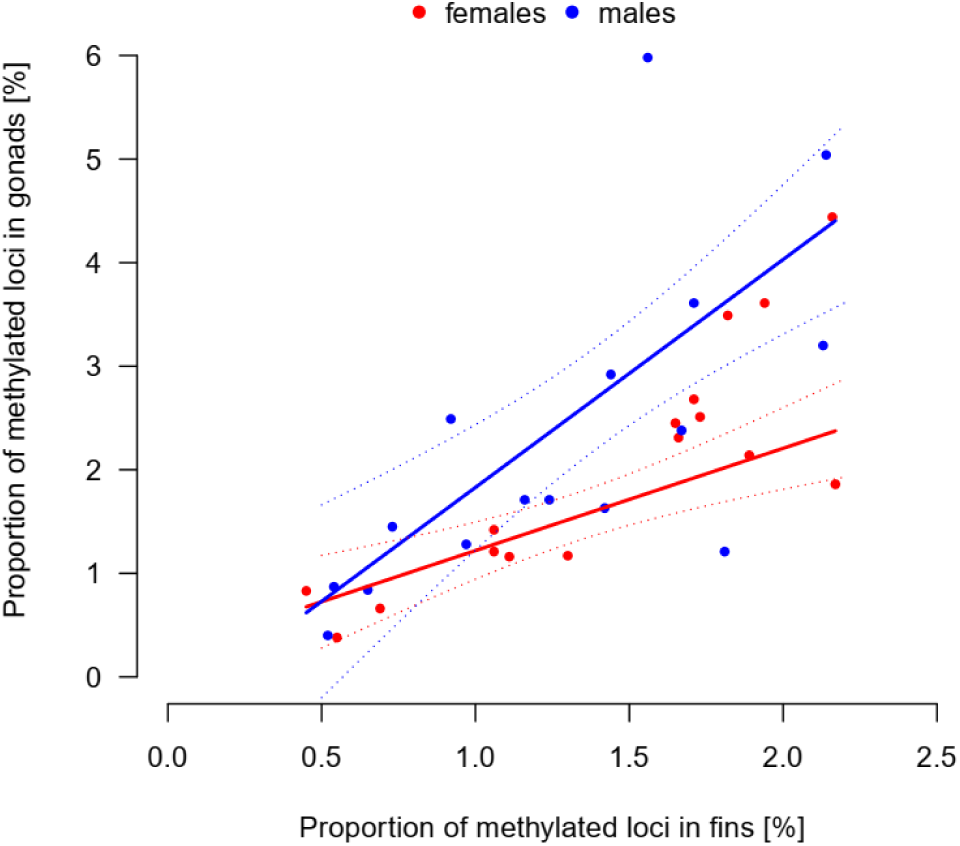
Positive relationship between the proportion of methylated loci in the fins and gonads of the sampled individuals. Proportion of DNA methylation in gonads increased more steeply with the proportion of methylated loci in fins in males (blue points and trendline) than in females (red points and trendline). The dotted lines represent limits of 95% CI of the corresponding trendlines.

### Similarity of DNA methylation pattern between tissues

As shown by clustering analysis, the status of DNA methylation across the 1 296 differentially methylated loci (i.e., loci that were not uniformly methylated or unmethylated across all 64 samples) was very similar across fin samples (Fig. 6, all empty points). The similarity analysis showed that ovaries also clustered relatively closely together (Fig. 6, red full points), while male gonads were the most diverse (Fig. 6, blue full points), spreading across both the first and second axes of variation. Based on the DNA methylation patterns, the samples did not group meaningfully with respect to their age (see Figure S3).

**Figure 6.**
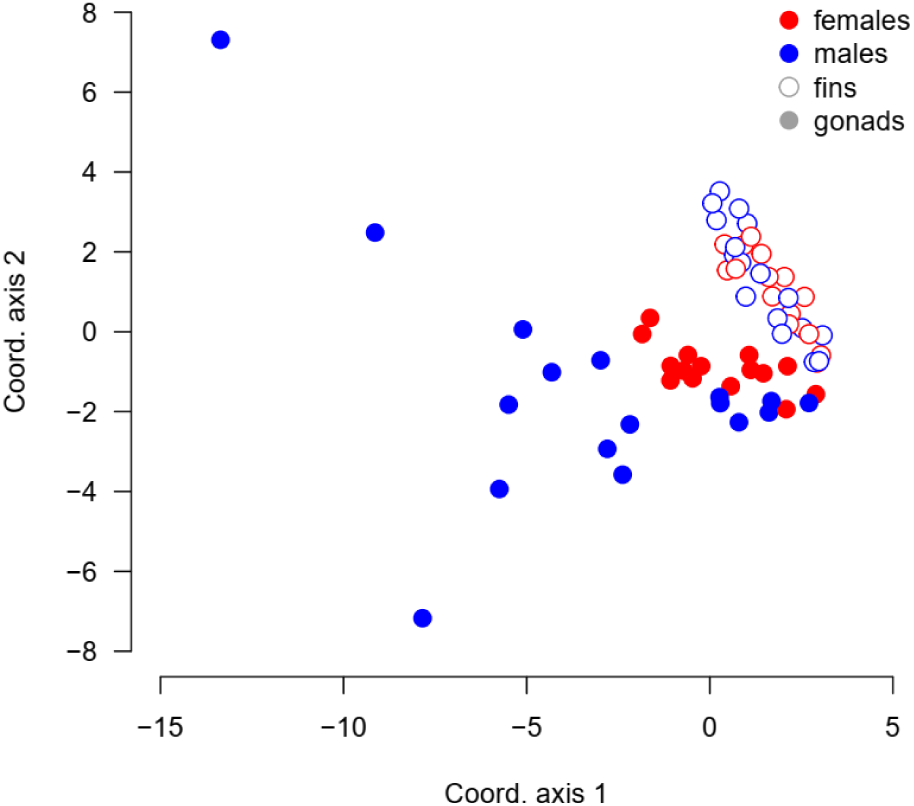
Ordination of DNA methylation similarity across samples based on classical multi- dimensional scaling. The samples group with regard to the tissue (fins – empty points, gonads – full points), and in the gonads with regard to sex (ovaries - red full points, testes – blue full points). Coordinate axis 1 and 2 explain 9.8% and 4.7% of the total variation in methylation status across samples, respectively.

### Shared methylated loci in fins and gonads across individuals

The similarity and clustering of samples from the same tissue can be best explained by shared unmethylated (not by shared methylated) loci. For fins of both sexes, the shared unmethylated pattern across samples represented a high proportion of the differentially methylated loci (808 for males and 843 for females, 65.0%). Similarly, large part of the differentially methylated loci was unmethylated across female gonads (718 loci, 55.4%). However, it was only 117 (8.8%) loci uniformly unmethylated across the gonads of different males. Looking at the uniformly methylated loci, we could not find any locus shared across all male or female fins or male gonads. There was only a single locus methylated across all female gonads.

Despite the high correlation in proportion of DNA methylation between fin and gonad, we did not find any locus with shared fin-gonad methylation status across all individuals. In the end, 8 to 88 of the differentially methylated loci (i.e., 0.62 to 6.79%) were methylated in both fins and gonads of the same individual. Not a single locus was methylated in both fins and gonads across all females (N = 15) or males (N = 17). This suggests that there was a large individual variation in the DNA methylation patterns.

## DISCUSSION

We compared the age-related dynamics of telomere length and DNA methylation between somatic and gonad tissue in a cross-sectional study of the turquoise killifish. Both telomere length and proportion of DNA methylation in fins corresponded well with the state in the gonads of the same individual. We recorded relatively low overall rate of methylated DNA and, despite the strong correlation, only few loci shared their methylation status between fins and gonads. In general, we demonstrate it is legitimate to use fins as a surrogate tissue for age-related changes of telomere length and the levels of DNA methylation in the gonads of the turquoise killifish.

Our main goal was to test for utility of fin sample as a proxy for age-related changes of telomere length and DNA methylation in fish gonad. To properly capture ageing dynamics, individuals should be sampled repeatedly during their lifetime as senescence trajectories can vary within the same population (Hamel et al., 2018; McCleery et al., 2008; Nussey et al., 2008). Repeated sampling is nevertheless difficult or impossible when targeting age-related changes of internal organs. One of the workarounds is using a reliable proxy tissue. Here, we propose using sample from non-regenerated part of fin to look at the status of telomeres and DNA methylation in gonads of the turquoise killifish, an emerging model organism for the study of ageing and senescence. We targeted fin biopsies because alternative sampling methods such as collecting blood would be too intrusive and potentially lethal to such a small vertebrate. Skin swabs or environmental DNA, on the other hand, are not sufficient for obtaining a reliable amount of good quality DNA for TRF analysis or ddRAD sequencing. In this sense, fin biopsy represents a relatively benign sampling method with satisfactory DNA yield. The impact of fin sampling on fish life is further mitigated by their high regeneration capacity (Mayne et al., 2021). Our promising results thus suggest that a similar approach should be tested in other small model fish species as well.

Rate of telomere shortening can vary dramatically between different organs (Cherif et al., 2003; Ramos-Ibeas et al., 2019). In small freshwater fish, the ageing dynamics of telomere lengths between soma and gonads appears to be highly variable (Carneiro et al., 2016; Lunghi and Bilandžija, 2024; Morbiato et al., 2023). We approached the issue directly and sampled three age groups specifically for fins and gonads in both males and females. Our results are supported by the use of the terminal restriction fragment (TRF) method, which is widely regarded as the gold standard for telomere measurement and is considerably less affected by stochastic variation than the frequently used qPCR (Kärkkäinen et al., 2022). In this respect, we believe we brought a robust support for the use of fin telomere lengths as a proxy for the state in ageing gonads of both sexes in the turquoise killifish.

Age-related changes of DNA methylation across multiple tissues have been largely understudied in fish models (for mammals see Kawamura et al., 2025; Lu et al., 2023; Spiers et al., 2016). A recent analysis of DNA methylation patterns in turquoise killifish fins and brain demonstrated large overlap in the differentially methylated regions and their status between these two functionally and physiologically divergent tissues (Giannuzzi et al., 2024). We examined both the number of methylated loci and their shared pattern across samples in fins and gonads. Fins appear to be a reliable indicator for the proportion of DNA methylation in gonads of the same individual, although one needs to account for the sex-specific differences. However, we did not identify any general commonly methylated loci across our samples. The agreement in loci methylation status in fin and gonad of the same individual was relatively low (up to 7%). This probably stems from the reduced representation sequencing method. The EpiRAD version of the ddRAD sequencing (Dimond et al., 2017; Schield et al., 2016) was adopted to estimate levels of DNA methylation levels across samples (elaborated in detail in Dianat et al., in preparation). Although ddRAD sequencing is relatively inexpensive and targeted to specific restriction sites, it typically covers only about 1% of the genome. As a result, a substantial proportion of shared methylation patterns may have been missed. Further focused study of the DNA methylation patterns across different tissues in the turquoise killifish is thus warranted.

We captured relatively low proportion of DNA methylation (maximum 6%; compared to 65- 80% in other teleosts as pufferfish, tilapia or zebrafish; Mayne et al., 2020; Wan et al., 2016; Zemach et al., 2010). The detected methylation thus represents only a small fraction of the overall methylation landscape. The low percentage of recovered DNA methylation results from the EpiRAD method itself and also from processing the sequencing data. The principle of the EpiRAD sequencing method (see e.g. Dimond et al., 2017) is obtaining two libraries from the same sample (once digested with methylation non-sensitive restriction enzyme and the other aliquot with a methylation sensitive equivalent). Compared to bisulfite sequencing, for example, the approach is indirect as we worked with two libraries from a single sample where one served as a template of the present genetic variation (RAD library) and the other (EpiRAD library) for quantification of which of the present loci are methylated. We then decided to identify methylation based only on the presence or absence of the specific locus in the methylation-sensitive library (EpiRAD library). With this approach, we avoid false positives of methylated loci, but also omit loci that were only partially methylated (represented by low number of reads). Searching for a threshold separating truly methylated and non-methylated loci can be difficult, because such a threshold would have to be sample specific. In the EpiRAD (methylation-sensitive) libraries, higher occurrence of methylated loci results in more reads from the non-methylated loci. The number of reads in a specific locus thus depends on the overall rate of DNA methylation in the sample and the threshold should account for that to recover true locus status. Applying uniform threshold across samples would result in underestimating DNA methylation proportion in relatively more methylated samples (with relatively less loci in the EpiRAD library), because the overall number of reads is concentrated among fewer loci and each locus therefore has relatively high number of reads. Less methylated samples (with more loci in the EpiRAD library), on the other hand, would be overestimated as the number of reads of some of the non-methylated loci would remain below the general threshold. There is currently no widely accepted statistical approach to analyse this type of data (i.e. general model to estimate level of methylation or probability of methylation from comparison of individual EpiRAD and RAD library pairs). Choosing a general arbitrary threshold for all samples is problematic and we decided to employ the most straightforward method - setting the general threshold to zero. The statistical framework for processing for the EpiRAD approach for epigenetics from compositional data needs to be further developed and validated with methods that identify DNA methylation directly.

The secondary aim of our work was to test for the changes related to increased age. We expected a continuous decline in telomere lengths in older AGE-groups (Hartmann et al., 2009) and higher proportion of methylated loci. While telomeres were shorter in the middle age-group (22 weeks after hatching) compared to the young fish (8 weeks old), old fish (at 30 weeks) had relatively similar telomere length with the young individuals. This concave pattern was probably caused by combined effects of survivor bias and random selection for sampling of small number of individuals. Inter-individual variability in telomere lengths (Figure S2) can then override possible age-related telomere shortening. While it may be possible that telomere attrition is not apparent with age under some circumstances or even telomere elongation could be observed (Anchelin et al., 2011; Morbiato et al., 2023), the earlier work by Hartmann et al. (2009) and our unpublished longitudinal data (Vrtilek et al., unpublished data) indeed demonstrate consistent yet subtle telomere shortening throughout the life of the turquoise killifish. The effect of age on proportion of methylated DNA was apparent in fins, but not in gonads of the turquoise killifish. DNA methylation in gonads showed relatively high individual variation, especially in males (Fig. 4B, Fig. 6). Age-associated increase of DNA methylation accumulating in the fin are in contrast with previous findings in zebrafish where muscle, liver, and brain tissues showed progressive demethylation with age (Shimoda et al., 2014). The high individual variability in gonads can come from germline cells often exhibiting more stable methylation patterns (Ortega-Recalde et al., 2019).

A potential shortcoming of our study is that we did not separate germline from the somatic cells of the sampled gonads. Germline composes only part of the gonad and is supported by somatic cells (e.g., Liu et al., 2022; Moses et al., 2024; Sposato et al., 2024). Extracting solely germline is difficult and obtaining sufficient DNA from oocytes, for example, is especially challenging due to the high requirement for the mass of the material (Monaghan and Metcalfe, 2019). Sampling sperm directly is more straightforward (Cattelan and Valenzano, 2023) but can introduce another bias due to selective survival of spermatozoa with longer telomeres, for example (Ramos-Ibeas et al., 2019; Sanghvi et al., 2024). We therefore focused on the whole gonads and this needs to be kept in mind. The presence of somatic cells in ovaries may partially explain their higher similarity with fins in the clustering analysis compared with testes that were relatively more dispersed. This may have even improved the fit of the fin-gonad correlations but given the relationship was strong for fin and both ovaries and testes, this inflation was likely minor.

## CONCLUSION

We gathered important methodological and biological insights that may advance ageing research in small laboratory fish. By combining telomere length measurements with assessments of DNA methylation levels, fins were demonstrated to serve as a suitable proxy tissue to control for age-related changes in the gonads of both female and male turquoise killifish. Future studies should focus on qualitative age-associated changes in DNA methylation within somatic and gonadal (or germline) tissues, which were beyond the scope of the present work.

## Supporting information

Full supplement information

## AUTHOR CONTRIBUTIONS

**Milan Vrtílek** – Conceptualization, Funding acquisition, Investigation, Data curation, Formal analysis, Visualization, Writing – original draft, Writing – review and editing

**Anna Kromerová** – Investigation, Data curation, Formal analysis, Visualization, Writing – review and editing

**Malahat Dianat** – Investigation, Data curation, Formal analysis, Writing – review and editing **Miloslava Fojtová** – Conceptualization, Methodology, Supervision, Writing – review and editing **Dagmar Čížková** – Conceptualization, Methodology, Supervision, Writing – review and editing **Jiří Fajkus** - Conceptualization, Funding acquisition, Writing – review and editing

## ACKNOWLEDGEMENT

The work was conducted in accordance with a project nr. 31/2019 approved by the Expert committee for animal welfare of the Institute of Vertebrate Biology of the Czech Academy of Sciences. Computational resources were provided by the e-INFRA CZ project (ID:90254), supported by the Ministry of Education, Youth and Sports of the Czech Republic. This study was supported by the Czech Science Foundation (22-21198S).

## DATA AVAILABILITY STATEMENT

Data and the R script is available from FigShare DOI: 10.6084/m9.figshare.29987764

