## Supplementary material for "Fins as a reliable surrogate tissue for age-related changes of telomeres and DNA methylation in gonads of a short-lived fish": Full supplement information

**Table S1.** Overview of size and investment in reproduction in the experimental fish.

**Table S2.** The effect of changing the threshold for filtering loci with low number of reads on the results.

**Figure S1.** The effect of increasing filtering threshold for well-represented loci for analysis of DNA methylation.

**Figure S2.** Telomere lengths in fins and gonads of fish at the three AGE-samplings.

**Figure S3.** Ordination of DNA methylation similarity across samples based on classical multi-dimensional scaling coded with respect to AGE.

**Appendix 1.** Modified double digest restriction-site associated DNA (ddRAD) sequencing protocol for identification of DNA methylation.

**Appendix 2.** Parameter settings for assembling the libraries and concatenate the data using iPyrad.

**Table S1.** Overview of size and investment in reproduction in the experimental fish. Total body length of the fish was measured as the distance from the tip of the mouth to the end of the caudal fin. Gonado-Somatic Index (GSI) was calculated as gonad weight/dissected body weight).

| Sampling point | N | | Total body length [mm] | | Live body weight [g] | | GSI | |
|---|---|---|---|---|---|---|---|---|
| | Female | Male | Female | Male | Female | Male | Female | Male |
| AGE1 | 5 | 5 | 42.1 (±0.6) | 51.0 (±0.6) | 0.99 (±0.04) | 1.75 (±0.07) | 0.179 (±0.029) | 0.006 (±0.001) |
| AGE2 | 6 | 5 | 50.3 (±1.1) | 69.6 (±0.9) | 1.63 (±0.12) | 4.77 (±0.23) | 0.184 (±0.013) | 0.015 (±0.001) |
| AGE3 | 4 | 7 | 51.7 (±1.5) | 68.6 (±1.5) | 1.81 (±0.15) | 4.28 (±0.30) | 0.184 (±0.028) | 0.017 (±0.001) |

**Table S2.** The effect of changing the threshold for filtering loci with low number of reads on the results. All the RAD libraries had to have at least given number of reads ("Threshold") for the locus in question to keep it in the dataset. See also the effect of increasing the threshold from 0 to 100 on the resulting number of loci in RAD libraries and the proportion of methylation in Figure S1. The dataset based on setting the threshold to 15 (shaded row) was used in the downstream analyses. DNA methylation % was proportion of loci with zero reads in the methylation-sensitive EpiRAD library.

| Threshold | Total number of loci after filtering | Mean number of reads per locus after filtering | Mean % DNA methylation | Effect of AGE | Best model of DNA methylation in gonads on fins based on AICc, $R^2$ | Similarity ordination analysis |
|---|---|---|---|---|---|---|
| 0 | 16 701 | 49 | 7.12 | Fins P-value = 0.409<br>Gonads P-value = 0.242 | SEX*Fin DNAm interaction, 0.825 | Fins and gonads more separated |
| 7 | 9 571 | 75 | 2.42 | Fins P-value = 0.042<br>Gonads P-value = 0.091 | SEX*Fin DNAm interaction, 0.730 | Similar to 15 |
| 15 | 7 841 | 85 | 1.76 | Fins P-value = 0.019<br>Gonads P-value = 0.073 | SEX*Fin DNAm interaction, 0.689 | Fig. 6 |
| 21♣ | 7 104 | 89 | 1.58 | Fins P-value = 0.015<br>Gonads P-value = 0.082 | SEX*Fin DNAm interaction, 0.713 | Everything more similar than in 15 |
| 42 | 4 183 | 101 | 1.30 | Fins P-value = 0.005<br>Gonads P-value = 0.049 | SEX + Fin DNAm additive, 0.590 | Everything more similar than in 15 |

♣ The average number of reads per locus before filtering was 21

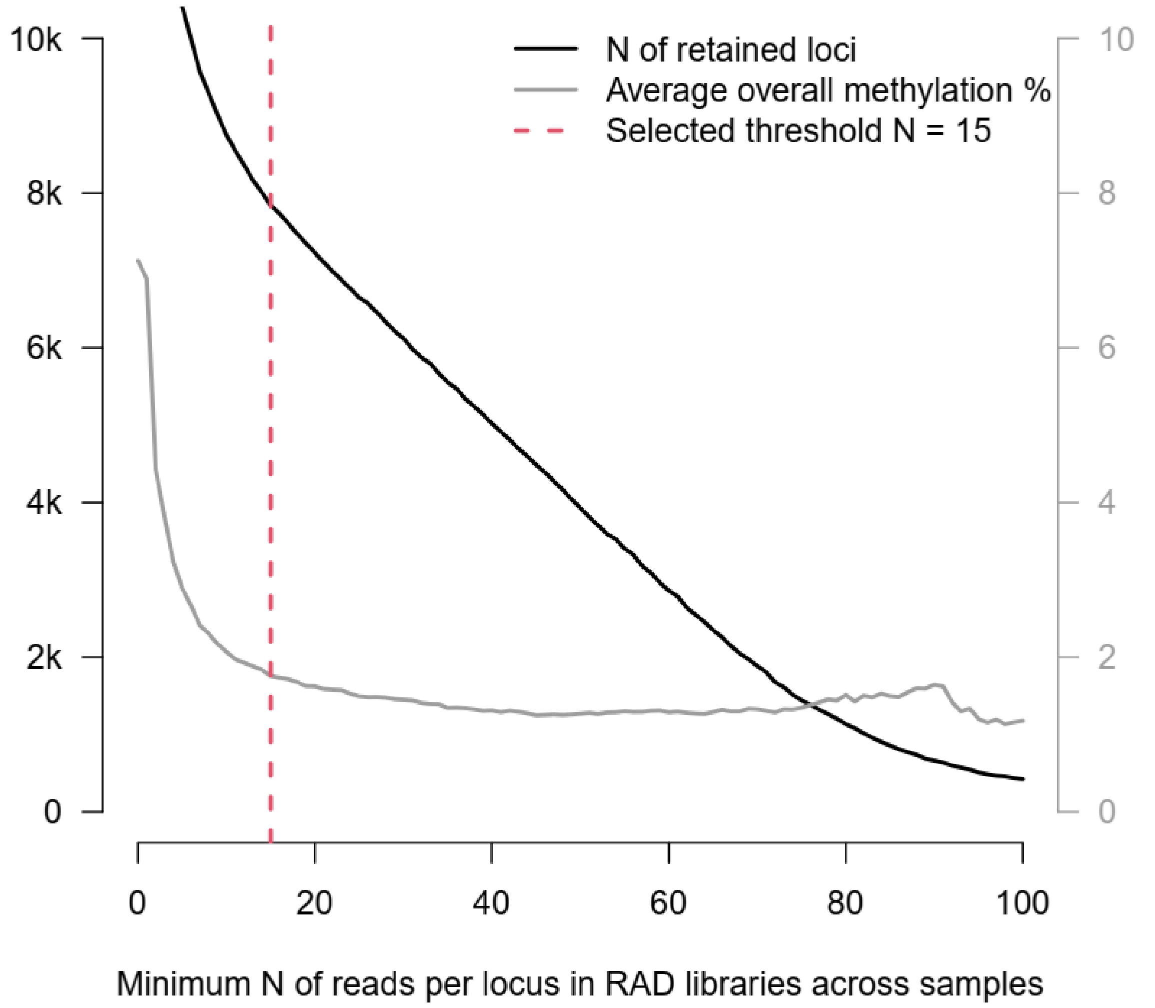


**Figure S1.** The effect of increasing filtering threshold for well-represented loci for analysis of DNA methylation. As the minimum read depth threshold per locus increases, the number of retained loci decreases. Additionally, the overall proportion of DNA methylation declines slightly before reaching a plateau. We decided to set the threshold to a minimum 15 reads per locus across all samples to retain sufficient number of loci for the analysis as the decline in DNA methylation proportion begins to stabilize.

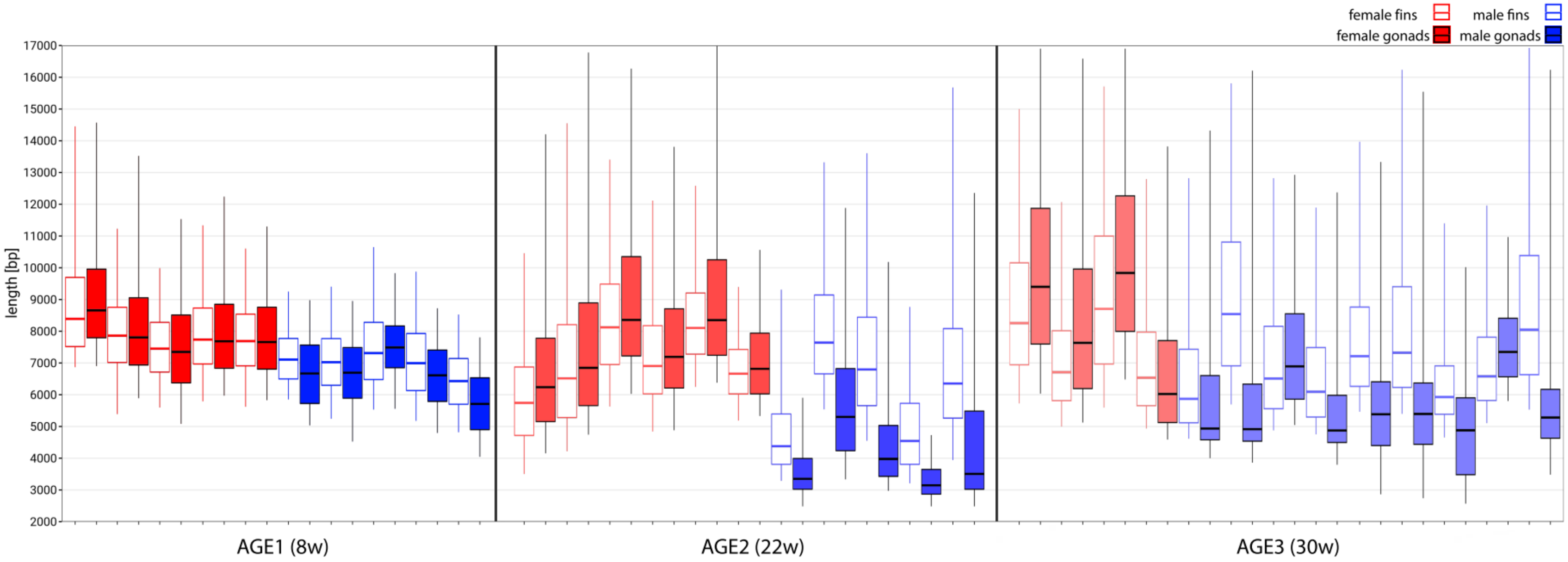


**Figure S2.** Telomere lengths in fins and gonads of fish at the three AGE-samplings. Lengths of telomeres were analysed by the TRF method, and telomere-specific hybridization signals were evaluated by the WALTER toolset (Lyčka et al., 2021). Top and bottom quartiles are separated by the median. Female, in red; male, in blue; fins, open boxes; gonads, filled boxes. Neighbouring samples (fin and gonad) were taken from the same fish.

(Lyčka, M., Peska, V., Demko, M., Spyroglou, I., Kilar, A., Fajkus, J., Fojtová, M., 2021. WALTER: an easy way to online evaluate telomere lengths from terminal restriction fragment analysis. BMC Bioinformatics 22, 1–14. https://doi.org/10.1186/s12859-021-04064-0)

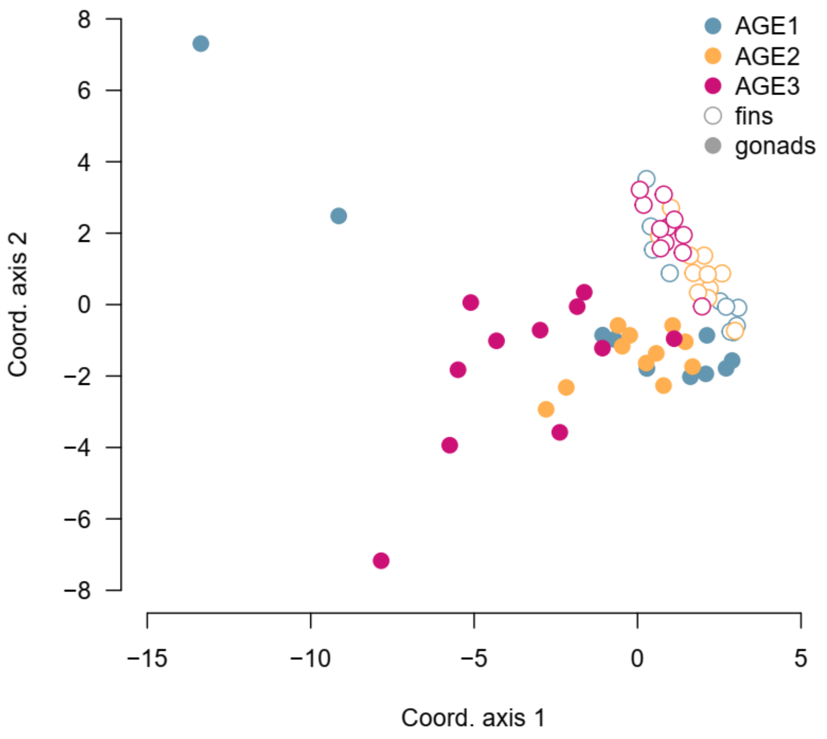


**Figure S3.** Ordination of DNA methylation similarity across samples based on classical multi-dimensional scaling coded with respect to AGE. The data is the same as for Figure 6 but colour-coded according to the AGE grouping. Coordinate axis 1 and 2 explain 9.8 % and 4.7 % of the total variation in methylation status across samples, respectively.

**Appendix 1.** Modified double digest restriction-site associated DNA (ddRAD-seq) protocol for identification of DNA methylation based on Peterson *et al*. (2012).

### DNA extraction

We extracted DNA from the turquoise killifish fin and gonad using the cetyltrimethylammonium bromide method (Kovařík et al., 2000). The DNA integrity was verified by standard horizontal agarose gel electrophoresis, and DNA concentration was measured by the NanoDrop spectrophotometer (Thermo Fisher Scientific) and Qubit (Thermo Fisher Scientific). We aimed to obtain DNA at a concentration of 20 ng/µL; however, we also included samples with lower concentration. For ddRAD library preparation, we prioritized processing DNA samples with relatively similar concentrations together**.**

### Adapter preparation

Oligos (Table 1) were initially eluted in TLE buffer to a final concentration of 100 µM (stock). To prepare the adapters, 50 µL of each oligo from a pair were mixed and went through the following protocol in a thermal cycler: 95°C for 1 min, followed by slow cooling of -0.1°C per second to 20°C. The i5 adapters were then diluted in TLE to a working concentration of 10 µM. Finally, they were aliquoted and stored at -20°C.

Table 1. Oligo pairs used to prepare adapters. * = PTO-Phosphorothioates, [PHO] = 5´phosphorylation. Inline barcode is in blue font.

| | |
|---|---|
| i5-upper1 | ACGACGCTCTTCCGATCTCATCC*A |
| i5-lower1 | [PHO]AATTTGGATGAGATCGGAAGAGCGTCGTGTAGGGAAAGAGTGT |
| i5-upper2 | ACGACGCTCTTCCGATCTAGCAAT*C |
| i5-lower2 | [PHO]AATTGATTGCTAGATCGGAAGAGCGTCGTGTAGGGAAAGAGTGT |
| i7-upper1 | [PHO]CGAGATCGGAAGAGCACACGTaatcc |
| i7-lower1 | GTGACTGGAGTTCAGACGTGTGCTCTTCCGATC*T |
| i7-upper2 | [PHO]CGTAGATCGGAAGAGCACACGTaatcc |
| i7-lower2 | GTGACTGGAGTTCAGACGTGTGCTCTTCCGATCT*A |

### Protocol for ddRAD sequencing:

#### 1) Restriction reaction:

REAGENT 1 and REAGENT 2 restriction mixes (Table 2) were incubated at 37 °C for 4 hours.

Table 2. Reagents for the restriction reactions.

| REAGENT 1, non-sensitive to methylation | Per sample (µL) |
|---|---|
| $H_2O$ | 6 |
| EcoRI-HF restriction enzyme (20,000 U/ml) | 0.5 |
| MspI restriction enzyme (20,000 U/ml) | 0.5 |
| CutSMART buffer (10× concentrated) | 3 |
| Genomic DNA | 20 |
| TOTAL VOLUME | 30 |

| REAGENT 2, sensitive to methylation | Per sample (µL) |
|---|---|
| $H_2O$ | 5.5 |
| EcoRI-HF restriction enzyme (20,000 U/ml) | 0.5 |
| HpaII restriction enzyme (5000 U, 10U/µL) | 1 |
| CutSMART buffer (10× concentrated) | 3 |
| Genomic DNA | 20 |
| TOTAL VOLUME | 30 |

### 2) Adapter ligation:

During ligation, the DNA fragments were attached to i5 and i7 adapters consisting of inline barcodes (only at i5) (Table 1). Ligation (Table 3) was done using this protocol: 23 °C for 30 minutes, 65 °C for 10 minutes, then the temperature decreased to 22 °C by 2 °C/cycle.

Table 3. Reagents for the ligation reaction.

| REAGENT | Per sample (µL) |
|---|---|
| T4 DNA ligase (400 kU/ml) | 0.1 |
| Buffer (10× concentrated) | 4 |
| i5 adapter (10 µM) | 0.8 |
| i7 adapter (100 µM) | 1.2 |
| Digested DNA | 17 |
| $H_2O$ | 16.9 |

### 3) SpriSelect cleaning 1x

We followed the manufacturer's protocol for cleaning up the resulting 40 µL of DNA libraries with 40 µL of beads (1×). DNA was eluted by 20 µL of Tris buffer (10 mM); 10 µL used for PCR, 10 µL was stored as a backup.

#### 4) PCR amplification:

Indexes were added to the libraries during the PCR amplification step (Table 4). PCR was run under these conditions: 98 °C for 45 s; 12-17 cycles (depending on DNA concentration) of 98 °C for 15 s, 60 °C for 30 s, 72 °C for 30 s; and a final extension at 72 °C for 1 min.

Table 4. Reagents for the PCR.

| REAGENT | Per sample (µL) |
|---|---|
| iTru primer p5 (indexed) (10 µM) | 1.25 |
| iTru primer p7 (indexed) (10 µM) | 1.25 |
| KAPA HiFi HS Ready mix | 12.5 |
| Adapter-ligated library | 10 |
| Total volume | 25 |

We used unique combinations (per sample) of 0.5 µM iTrue primers (Glenn et al., 2019), resulting in a dual indexed sequencing library (Table 5). Proper amplification was tested by agarose gel electrophoresis using 5 µL of PCR product.

Table 5. Indexed iTru p5 and p7 primers (Glenn et al., 2019).

| PRIMER NAME | Primer sequence |
|---|---|
| iTru5_03_A | AATGATACGGCGACCACCGAGATCTACACAACACCACACACTCTTTCCCTA*C |
| iTru5_03_B | AATGATACGGCGACCACCGAGATCTACACTGAGCTGTACACTCTTTCCCTA*C |
| iTru5_03_C | AATGATACGGCGACCACCGAGATCTACACCACAGGAAACACTCTTTCCCTA*C |
| iTru5_03_D | AATGATACGGCGACCACCGAGATCTACACTGACAACCACACTCTTTCCCTA*C |
| iTru5_03_E | AATGATACGGCGACCACCGAGATCTACACTGTTCCGTACACTCTTTCCCTA*C |
| iTru5_03_F | AATGATACGGCGACCACCGAGATCTACACCCTAGAGAACACTCTTTCCCTA*C |
| iTru5_03_G | AATGATACGGCGACCACCGAGATCTACACGCATAACGACACTCTTTCCCTA*C |
| iTru5_03_H | AATGATACGGCGACCACCGAGATCTACACCAGTGCTTACACTCTTTCCCTA*C |
| iTru5_05_A | AATGATACGGCGACCACCGAGATCTACACGGTACGAAACACTCTTTCCCTA*C |
| iTru5_05_B | AATGATACGGCGACCACCGAGATCTACACAAGCATCGACACTCTTTCCCTA*C |
| iTru5_05_C | AATGATACGGCGACCACCGAGATCTACACGCCAATACACACTCTTTCCCTA*C |
| iTru5_05_D | AATGATACGGCGACCACCGAGATCTACACCTGTATGCACACTCTTTCCCTA*C |
| iTru5_05_E | AATGATACGGCGACCACCGAGATCTACACCTTAGGACACACTCTTTCCCTA*C |
| iTru5_05_F | AATGATACGGCGACCACCGAGATCTACACTCAGCCTTACACTCTTTCCCTA*C |
| iTru5_05_G | AATGATACGGCGACCACCGAGATCTACACACATGCCAACACTCTTTCCCTA*C |
| iTru5_05_H | AATGATACGGCGACCACCGAGATCTACACGATGGAGTACACTCTTTCCCTA*C |
| iTru7_201_10 | CAAGCAGAAGACGGCATACGAGATTGGTATCCGTGACTGGAGTTCA*G |
| iTru7_201_11 | CAAGCAGAAGACGGCATACGAGATGATGTCGAGTGACTGGAGTTCA*G |
| iTru7_204_10 | CAAGCAGAAGACGGCATACGAGATGAGAAGGTGTGACTGGAGTTCA*G |
| iTru7_204_11 | CAAGCAGAAGACGGCATACGAGATTCTTACGGGTGACTGGAGTTCA*G |

| | |
|---|---|
| iTru7_205_10 | CAAGCAGAAGACGGCATACGAGATAACAGGTGGTGACTGGAGTTCA*G |
| iTru7_206_01 | CAAGCAGAAGACGGCATACGAGATAGTCTTGGGTGACTGGAGTTCA*G |
| iTru7_206_09 | CAAGCAGAAGACGGCATACGAGATTGTCAGTGGTGACTGGAGTTCA*G |
| iTru7_206_11 | CAAGCAGAAGACGGCATACGAGATACAGTGACGTGACTGGAGTTCA*G |
| iTru7_206_12 | CAAGCAGAAGACGGCATACGAGATGTTGGCATGTGACTGGAGTTCA*G |
| iTru7_208_03 | CAAGCAGAAGACGGCATACGAGATTGCAAGACGTGACTGGAGTTCA*G |
| iTru7_210_03 | CAAGCAGAAGACGGCATACGAGATTCACTCGAGTGACTGGAGTTCA*G |
| iTru7_401_08 | CAAGCAGAAGACGGCATACGAGATAACCTACGGTGACTGGAGTTCA*G |
| iTru7_201_09 | CAAGCAGAAGACGGCATACGAGATCAACCGTAGTGACTGGAGTTCA*G |
| iTru7_202_07 | CAAGCAGAAGACGGCATACGAGATAGAGACTCGTGACTGGAGTTCA*G |
| iTru7_202_12 | CAAGCAGAAGACGGCATACGAGATCACGCAATGTGACTGGAGTTCA*G |
| iTru7_203_05 | CAAGCAGAAGACGGCATACGAGATTGGTTCGAGTGACTGGAGTTCA*G |
| iTru7_204_06 | CAAGCAGAAGACGGCATACGAGATATCCGTTGGTGACTGGAGTTCA*G |
| iTru7_206_03 | CAAGCAGAAGACGGCATACGAGATACACTACCGTGACTGGAGTTCA*G |
| iTru7_208_04 | CAAGCAGAAGACGGCATACGAGATTGCACTTGGTGACTGGAGTTCA*G |
| iTru7_208_12 | CAAGCAGAAGACGGCATACGAGATAGTTCGCAGTGACTGGAGTTCA*G |
| iTru7_210_06 | CAAGCAGAAGACGGCATACGAGATTCACCTAGGTGACTGGAGTTCA*G |
| iTru7_401_01 | CAAGCAGAAGACGGCATACGAGATTCGTGCATGTGACTGGAGTTCA*G |
| iTru7_401_02 | CAAGCAGAAGACGGCATACGAGATATGGCGATGTGACTGGAGTTCA*G |
| iTru7_402_11 | CAAGCAGAAGACGGCATACGAGATTGCTTGCTGTGACTGGAGTTCA*G |

#### 5) Measurement of PCR product concentration:

The approximate concentrations of PCR products were estimated using agarose gel electrophoresis according to the intensities of bands of the 100 bp ladder (Invitrogen, Thermo Fisher Scientific, USA).

#### 6) Pooling and re-concentration of samples:

We pooled the samples equimolarly based on their concentrations, creating one subpool per gel. These subpools were subsequently pooled equimolarly into a final pool, which was then purified using SpriSelect (1.2×) and eluted in 55 µL of Tris buffer (10 mM).

We performed this step separately for the libraries derived from reagent 1 and reagent 2, and then combined the two libraries at the final stage.

#### 7) Size selection of library pool on Pippin-prep (Sage Science, USA):

Size selection was performed following the manufacturer's instructions. The libraries were size-selected in 306-340-374 bp.

#### 8) Sequencing:

The final library was sent for sequencing on the Illumina NovaSeq X Plus PE150 platform at Novogene Co. Ltd., obtaining 150 bp paired-end (PE) reads.

**Appendix 2.** Parameter settings for assembling the libraries and concatenate the data using iPyrad.

### RAD libraries

```
notho_msp_PSE3                  ## [0] [assembly_name]: Assembly name. Used to name output directories for assembly steps
./                              ## [1] [project_dir]: Project dir (made in curdir if not present)
                                ## [2] [raw_fastq_path]: Location of raw non-demultiplexed fastq files
                                ## [3] [barcodes_path]: Location of barcodes file
./../*fastq                     ## [4] [sorted_fastq_path]: Location of demultiplexed/sorted fastq files
reference                       ## [5] [assembly_method]: Assembly method (denovo, reference, denovo+reference, denovo-reference)
./../GCF_027789165.1_UI_Nfuz_MZM_1.0_genomic.fna  ## [6] [reference_sequence]: Location of reference sequence file
pairddrad                       ## [7] [datatype]: Datatype (see docs): rad, gbs, ddrad, etc.
AATTC, CGG                      ## [8] [restriction_overhang]: Restriction overhang (cut1,) or (cut1, cut2)
5                               ## [9] [max_low_qual_bases]: Max low quality base calls (Q<20) in a read
33                              ## [10] [phred_Qscore_offset]: phred Q score offset (33 is default and very standard)
6                               ## [11] [mindepth_statistical]: Min depth for statistical base calling
6                               ## [12] [mindepth_majrule]: Min depth for majority-rule base calling
10000                           ## [13] [maxdepth]: Max cluster depth within samples
0.9                             ## [14] [clust_threshold]: Clustering threshold for de novo assembly
0                               ## [15] [max_barcode_mismatch]: Max number of allowable mismatches in barcodes
2                               ## [16] [filter_adapters]: Filter for adapters/primers (1 or 2=stricter)
35                              ## [17] [filter_min_trim_len]: Min length of reads after adapter trim
2                               ## [18] [max_alleles_consens]: Max alleles per site in consensus sequences
0.05                            ## [19] [max_Ns_consens]: Max N's (uncalled bases) in consensus (R1, R2)
0.08                            ## [20] [max_Hs_consens]: Max Hs (heterozygotes) in consensus (R1, R2)
2                               ## [21] [min_samples_locus]: Min # samples per locus for output
0.2                             ## [22] [max_SNPs_locus]: Max # SNPs per locus (R1, R2)
8                               ## [23] [max_Indels_locus]: Max # of indels per locus (R1, R2)
0.5                             ## [24] [max_shared_Hs_locus]: Max # heterozygous sites per locus (R1, R2)
0, 0, 0, 0                      ## [25] [trim_reads]: Trim raw read edges (R1>, <R1, R2>, <R2) (see docs)
0, 0, 0, 0                      ## [26] [trim_loci]: Trim locus edges (see docs) (R1>, <R1, R2>, <R2)
*                               ## [27] [output_formats]: Output formats (see docs)
                                ## [28] [pop_assign_file]: Path to population assignment file
```

### EpiRAD libraries

```
notho_hpa_PSE3                  ## [0] [assembly_name]: Assembly name. Used to name output directories for assembly steps
./                              ## [1] [project_dir]: Project dir (made in curdir if not present)
                                ## [2] [raw_fastq_path]: Location of raw non-demultiplexed fastq files
                                ## [3] [barcodes_path]: Location of barcodes file
./../*fastq                     ## [4] [sorted_fastq_path]: Location of demultiplexed/sorted fastq files
reference                       ## [5] [assembly_method]: Assembly method (denovo, reference, denovo+reference, denovo-reference)
```

```
./../GCF_027789165.1_UI_Nfuz_MZM_1.0_genomic.fna  ## [6] [reference_sequence]: Location of reference sequence file
pairddrad                      ## [7] [datatype]: Datatype (see docs): rad, gbs, ddrad, etc.
AATTC, CGG                     ## [8] [restriction_overhang]: Restriction overhang (cut1,) or (cut1, cut2)
5                              ## [9] [max_low_qual_bases]: Max low quality base calls (Q<20) in a read
33                             ## [10] [phred_Qscore_offset]: phred Q score offset (33 is default and very standard)
6                              ## [11] [mindepth_statistical]: Min depth for statistical base calling
6                              ## [12] [mindepth_majrule]: Min depth for majority-rule base calling
10000                          ## [13] [maxdepth]: Max cluster depth within samples
0.9                            ## [14] [clust_threshold]: Clustering threshold for de novo assembly
0                              ## [15] [max_barcode_mismatch]: Max number of allowable mismatches in barcodes
2                              ## [16] [filter_adapters]: Filter for adapters/primers (1 or 2=stricter)
35                             ## [17] [filter_min_trim_len]: Min length of reads after adapter trim
2                              ## [18] [max_alleles_consens]: Max alleles per site in consensus sequences
0.05                           ## [19] [max_Ns_consens]: Max N's (uncalled bases) in consensus (R1, R2)
0.08                           ## [20] [max_Hs_consens]: Max Hs (heterozygotes) in consensus (R1, R2)
2                              ## [21] [min_samples_locus]: Min # samples per locus for output
0.2                            ## [22] [max_SNPs_locus]: Max # SNPs per locus (R1, R2)
8                              ## [23] [max_Indels_locus]: Max # of indels per locus (R1, R2)
0.5                            ## [24] [max_shared_Hs_locus]: Max # heterozygous sites per locus (R1, R2)
0, 0, 0, 0                     ## [25] [trim_reads]: Trim raw read edges (R1>, <R1, R2>, <R2) (see docs)
0, 0, 0, 0                     ## [26] [trim_loci]: Trim locus edges (see docs) (R1>, <R1, R2>, <R2)
*                              ## [27] [output_formats]: Output formats (see docs)
                               ## [28] [pop_assign_file]: Path to population assignment file
```

### Concatenation

```
notho_cat_PSE3                 ## [0] [assembly_name]: Assembly name. Used to name output directories for assembly steps
./                             ## [1] [project_dir]: Project dir (made in curdir if not present)
                               ## [2] [raw_fastq_path]: Location of raw non-demultiplexed fastq files
                               ## [3] [barcodes_path]: Location of barcodes file
./../*fastq                    ## [4] [sorted_fastq_path]: Location of demultiplexed/sorted fastq files
reference                      ## [5] [assembly_method]: Assembly method (denovo, reference, denovo+reference, denovo-reference)
./../GCF_027789165.1_UI_Nfuz_MZM_1.0_genomic.fna  ## [6] [reference_sequence]: Location of reference sequence file
pairddrad                      ## [7] [datatype]: Datatype (see docs): rad, gbs, ddrad, etc.
AATTC, CGG                     ## [8] [restriction_overhang]: Restriction overhang (cut1,) or (cut1, cut2)
5                              ## [9] [max_low_qual_bases]: Max low quality base calls (Q<20) in a read
33                             ## [10] [phred_Qscore_offset]: phred Q score offset (33 is default and very standard)
6                              ## [11] [mindepth_statistical]: Min depth for statistical base calling
6                              ## [12] [mindepth_majrule]: Min depth for majority-rule base calling
10000                          ## [13] [maxdepth]: Max cluster depth within samples
0.9                            ## [14] [clust_threshold]: Clustering threshold for de novo assembly
```

```
0                              ## [15] [max_barcode_mismatch]: Max number of allowable mismatches in barcodes
2                              ## [16] [filter_adapters]: Filter for adapters/primers (1 or 2=stricter)
35                             ## [17] [filter_min_trim_len]: Min length of reads after adapter trim
2                              ## [18] [max_alleles_consens]: Max alleles per site in consensus sequences
0.05                           ## [19] [max_Ns_consens]: Max N's (uncalled bases) in consensus (R1, R2)
0.08                           ## [20] [max_Hs_consens]: Max Hs (heterozygotes) in consensus (R1, R2)
2                              ## [21] [min_samples_locus]: Min # samples per locus for output
0.2                            ## [22] [max_SNPs_locus]: Max # SNPs per locus (R1, R2)
8                              ## [23] [max_Indels_locus]: Max # of indels per locus (R1, R2)
0.5                            ## [24] [max_shared_Hs_locus]: Max # heterozygous sites per locus (R1, R2)
0, 0, 0, 0                     ## [25] [trim_reads]: Trim raw read edges (R1>, <R1, R2>, <R2) (see docs)
0, 0, 0, 0                     ## [26] [trim_loci]: Trim locus edges (see docs) (R1>, <R1, R2>, <R2)
*                              ## [27] [output_formats]: Output formats (see docs)
                               ## [28] [pop_assign_file]: Path to population assignment file
```
